# A Novel Gene Synthesis Platform for Designing Functional Protein Polymers

**DOI:** 10.1101/2024.09.01.610679

**Authors:** Toshimasa Homma, Rie Yamamoto, Lily Zuin Ping Ang, Alaa Fehaid, Mitsuhiro Ebara

## Abstract

Recombinant protein polymers with repeat sequences of specific amino acids can be regarded as sustainable functional materials that can be designed using genetic engineering. However, synthesizing genes encoding these proteins is significantly time-consuming and labor-intensive owing to the difficulty of using common gene synthesis tools, such as restriction enzymes and PCR primers. To overcome these obstacles, we propose a novel method: seamless cloning of rolling-circle amplicons (SCRCA). This method involves one-pot preparation of repetitive-sequence genes with overlapping ends for cloning, facilitating the easy construction of the desired recombinants. Using SCRCA, we synthesized 10 genes encoding hydrophilic resilin-like and hydrophobic elastin-like repeat units that induce liquid–liquid phase separation. SCRCA shows higher transformation efficiency and better workability than conventional methods, and the time and budget required for SCRCA are comparable to those required for non-repetitive-sequence gene synthesis. Additionally, SCRCA allows the construction of a repeat unit library at a low cost. The library shows considerably higher diversity compared with that of the state-of-the-art method. By combining this library construction with the directed evolution concept, we can rapidly develop an elastin-like protein polymer with a desired function. SCRCA can greatly accelerate research on protein polymers.

## 1. Introduction

Diverse repetitive amino acid sequences exist in nature, each of which has been found to possess distinctive functions.[1–5] Recombinant protein polymers inspired by these findings have attracted attention as sustainable and biocompatible materials synthesized using environment-friendly processes,[6,7] and the applications of these polymers are expanding to high-toughness fibers,[8,9] ultra-high elastic materials,[10,11] cell culture substrates,[12,13] bioseparation,[14,15] drug delivery,[16,17] proton-conducting membranes,[18] and smart cells.[19] The ability to freely design repeat units and chain lengths via genetic recombination technology represents a major advantage during the rational development of new protein-polymer materials.[3,20–22] Recently, copolymers and block copolymers composed of two repeat units with different functions have also been designed.[23–26]

To accelerate research in this field, the associated experimental operations must be streamlined. However, synthesis of repetitive-sequence genes encoding such proteins requires time-consuming methods because there are few unique parts whose sequences can be recognized using restriction enzymes or PCR primers.[27,28] Although a codon-scrambling algorithm has been developed to improve gene complexity,[29] screening for useful sequences requires expensive efforts. Rolling-circle amplification (RCA) is promising because it can simultaneously synthesize multiple repetitive-sequence genes with different repeat numbers.[30,31] However, the previously reported RCA methods are complicated to implement and unsuitable for the synthesis of diverse recombinants.

Our novel gene synthesis method, seamless cloning of rolling-circle amplicons (SCRCA), addresses this via four steps: preparation of the ssDNA ring template, RCA reaction, selecting the size of DNA fragments, and seamless cloning (**Figure 1a**). Seamless cloning simplifies the introduction of DNA fragments into a plasmid vector at the desired insertion position and in a specific direction, although overlapping sequences are required at both ends of the DNA fragments.[32] In SCRCA, these overlapping sequences are added at each end of the genes when the repeat sequence is synthesized, improving the success rate and versatility and significantly reducing the time and labor required for gene synthesis. By slightly modifying this method, copolymer and block polymer genes can also be synthesized using a single cloning operation (**Figure 1b** and **c**). Hence, SCRCA could contribute to the rapid and rational design of protein polymers.

**Figure 1.**
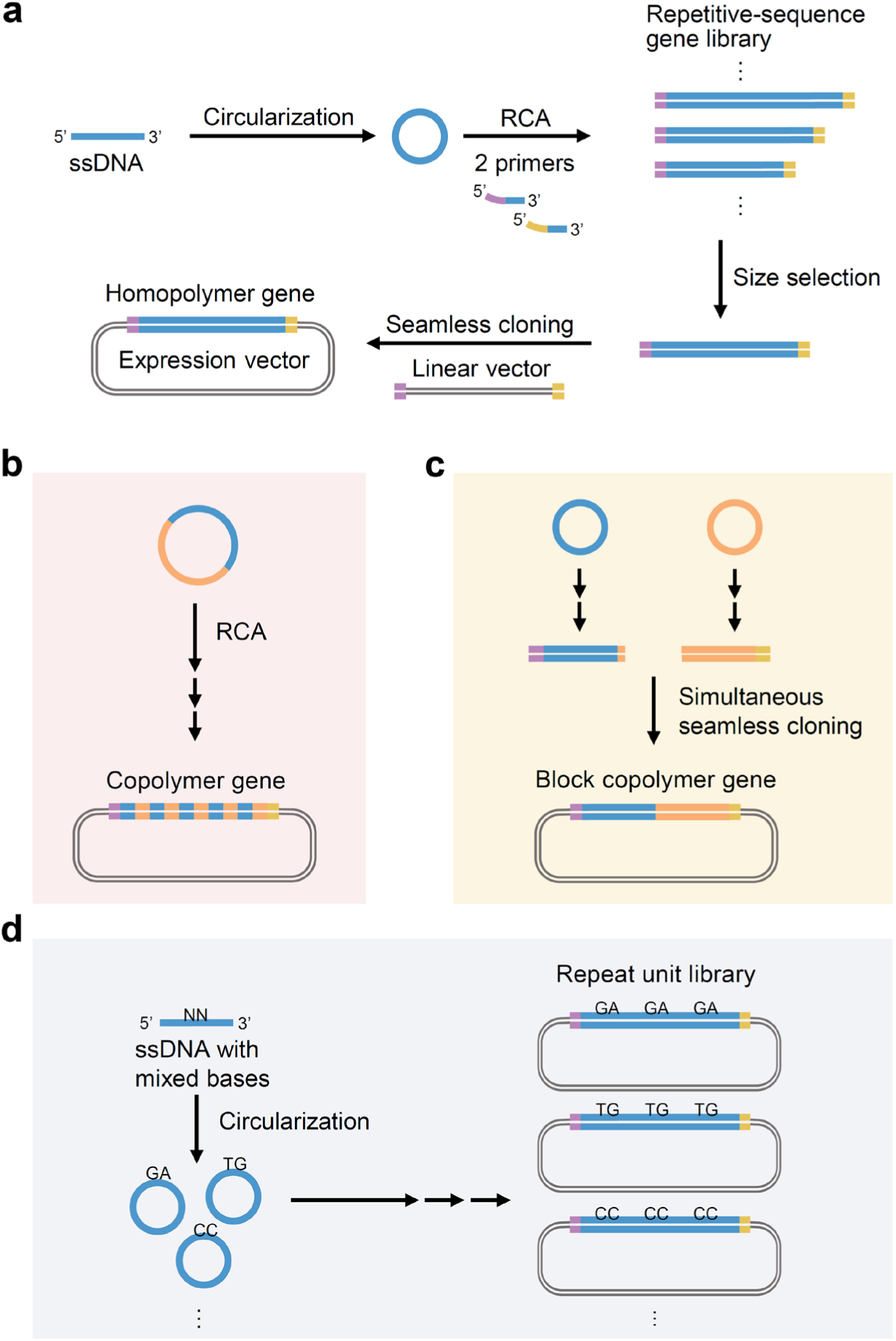
Seamless cloning of rolling-circle amplicons (SCRCA) (a) Synthesis of a homopolymer gene using the SCRCA method: 1) ssDNA oligo with a repeat unit sequence is circularized; 2) Repetitive-sequence gene library with different repeat numbers is prepared via RCA (blue line: repetitive sequence; purple and yellow lines: overlapping sequences); 3) Amplified products are separated based on their repeat numbers via agarose gel electrophoresis, and the gene of the desired length is excised; 4) This gene is inserted into an expression vector via seamless cloning. Copolymer and block copolymer genes composed of two different repeat units are synthesized via RCA using a large-ring template (b) and simultaneous seamless cloning (c), respectively. (d) Repeat unit library is constructed using an ssDNA oligo containing mixed bases.

In this study, we first synthesized repetitive-sequence genes encoding repeat units of a resilin-like polymer (RLP) and an elastin-like polymer (ELP), the major protein-polymer backbone, and demonstrated that the syntheses via SCRCA can be achieved with the same time and budget as the synthesis of non-repetitive sequence genes via common methods. Furthermore, we proved that repeat unit libraries with random mutations can be constructed at a low cost via SCRCA (**Figure 1d**). This facilitated the application of evolutionary molecular engineering to protein-polymer development. With the directed evolutionary experiment using the SCRCA-constructed libraries, we successfully developed a multi-responsive ELP in several months that would previously have required several years. These results confirm that SCRCA is a powerful tool for advancing protein polymer research.

## 2. Results and Discussion

### 2.1. Development of the Gene Amplification Process

For the SCRCA, we devised a new RCA reaction using forward and reverse primers with overlapping sequences at their 5′ ends. The Bst DNA polymerase large fragment was selected as the strand displacement enzyme for RCA because it lacks 5′→3′ exonuclease activity, and therefore, the overlapping sequence of the primer would not be digested.[33,34] Further, its high optimum temperature (60–70 °C)[35,36] is appropriate for amplifying GC-rich sequences, which are often present in major repetitive polypeptides. As an RCA template, a cyclized ssDNA with a sequence encoding the desired repeat unit was used. During isothermal DNA amplification using these materials, DNA elongation and strand displacement reactions are expected to occur continuously (Figure 2a). Consequently, a mixture of genes with repetitive sequences with different repeat numbers and overlapping sequences at both ends of the repetitive sequences can be prepared.

**Figure 2.**
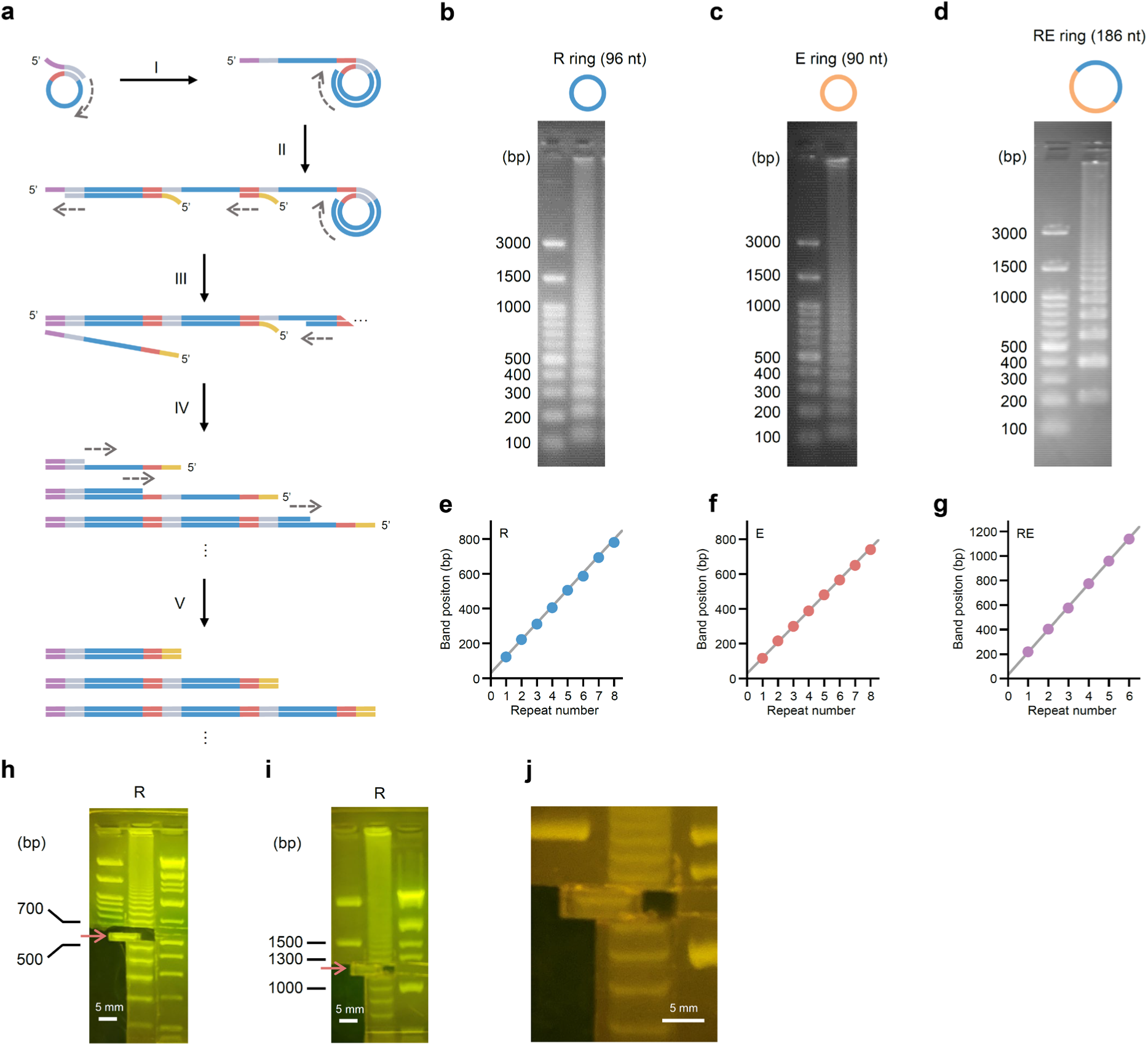
Isothermal rolling-circle amplification (RCA) for seamless cloning of rolling-circle amplicons (SCRCA) (a) Schematic of the isothermal amplification. Blue, gray, and red lines: repetitive sequences; purple and yellow lines: overlapping sequences. The 3′ ends of the forward and reverse primers are homologous to the gray and red lines, respectively. (I) A DNA chain, in which the complementary sequence of the ssDNA ring is repeated, is elongated from the reverse primer. (II) Forward primer attaches to the elongated DNA chain, forming a dsDNA section. (III) Nascent chain is dissociated by a subsequent elongation reaction. (IV) Primers with complementary sequences attach to the dissociated nascent chain to form a dsDNA fragment. (V) These reactions continue to occur to construct a library of repetitive genes with different repeat numbers. The 5′ and 3′ ends of the genes in the library have overlapping sequences derived from the 5′ ends of the forward and reverse primers, respectively. (b–d) Amplicons obtained by RCA using the (b) R, (c) E, and (d) RE rings were separated using 1.5% agarose gel electrophoresis. (e–g) Band positions of the amplicons obtained by RCA using the (e) R, (f) E, and (g) RE rings, and the linear function (gray line) of their theoretical lengths (repeat number × ring size + overlapping sequence bases) of these amplicons showed excellent correlation (R > 0.999). Each value represents the mean of three replicates, and the standard errors were less than 10 bp for all means. (h–j) Two DNA fragments equivalent to the 6 (h) and 12 repeats (i) of R ring sequences were cut from the gels electrophoresed for 70 and 110 mins, respectively. Panel (j) is an enlarged view of the cut gel, showing that the gel contained the 12-repeat DNA fragment.

To verify whether the reaction occurred as expected, isothermal amplification was performed using three ssDNA rings, namely R, E, and RE, that have nucleotide sequences encoding the hydrophilic RLP repeat unit[37] [(GRGDSPYS)_4_]_n_, hydrophobic ELP repeat unit[38] [(VGVPG)_6_]_n_, and RLP-ELP tandem repeat unit [GRGD-(SPYSGRGD)_3_-(GVPGVGVPGV)_6_-SPYS]_n_, respectively. RLP and ELP induce phase separation below an upper critical solution temperature and above a lower critical solution temperature (LCST), respectively. We considered that the advanced design of these polymers could contribute to the elucidation and application of the phase separation phenomenon.[39,40] Therefore, we adopted these polymers in our experimental models. Forward and reverse primers with 15-base overlapping sequences at their 5′ ends were used in the reactions. All reactions generated a mixture of amplified products of different lengths (Figure 2b**–d**). The positions of the bands matched the corresponding theoretical values (repeat number × ring size + overlapping sequence bases), indicating that the repetitive-sequence genes were successfully prepared (Figure 2e**–g**). As amplification products were obtained even when the large RE ring was used, SCRCA may also be suitable for gene synthesis of polymers with long repeat units and copolymers. As the amplified products could be separated by length using agarose gel electrophoresis, isolating the genes of the desired length was easy (Figure 2h). By increasing the running time of agarose gel electrophoresis, the length of the genes isolated could be increased (Figure 2i and j).

### 2.2. Repetitive-sequence Gene Construction

To evaluate the probability of constructing a transformant with the desired repetitive-sequence gene, *E. coli* transformation was performed using the In-Fusion seamless cloning system.[41] Homopolymer and copolymer genes *R6*, *E6*, *(RE)3*, *R12*, *E12*, and *(RE)6* were prepared using the RCA reactions using R, E, and RE rings (the repetitive-sequence genes are written in italics according to their repeat unit and repeat number; for example, *R6* represents a gene encoding a polypeptide with six repeats of the R ring sequence). Purified DNA fragments were subjected to fusion with the linear vector designed for expression and sequence confirmation (**Figure S1**). Block copolymer genes were synthesized via the simultaneous seamless cloning of two DNA fragments (Figure 1b). Here, four block copolymer genes, *R3E3*, *R8E4*, *R6E6*, and *R4E8* were prepared.

Colony PCR results showed that most of the transformants had the desired gene length (**Figure S2**). Sanger DNA sequencing revealed that 81.4% of the transformants contained the desired gene when the gene length was approximately 550 bp (*R6*, *E6*, *(RE)3*, and *R3E3*); even for genes with lengths >1000 bp (*R12*, *E12*, *(RE)6*, *R8E4*, *R6E6*, and *R4E8*), 41.5% of all the transformants exhibited the desired gene (Figure 3). Considering that a conventional method requires checking approximately 100 colonies to obtain a gene with a length >1000 bp;[30] this success rate is considerably high. Of the 32 sequences analyzed, there was one large deletion resulting from cloning failure and two deletions resulting from amplification of incomplete templates (**Figures S3–7**). Base substitution or deletion errors were less than 0.02% of >29,000 bases, comparable to those of the conventional RCA method.[30] Notably, only two weeks were required from ordering the oligo DNA to preparing the 10 repetitive-sequence genes. This duration is the same as that required for the outsourced synthesis of 1000-bp non-repetitive sequence genes. Furthermore, the cost of the DNA oligos and reagents required for the SCRCA was less than $500. Considering that the outsourced synthesis of four 500-bp and six 1000-bp non-repetitive sequence genes costs more than $2,000, the SCRCA method has solved the economic problem associated with repetitive-sequence gene synthesis.

**Figure 3.**
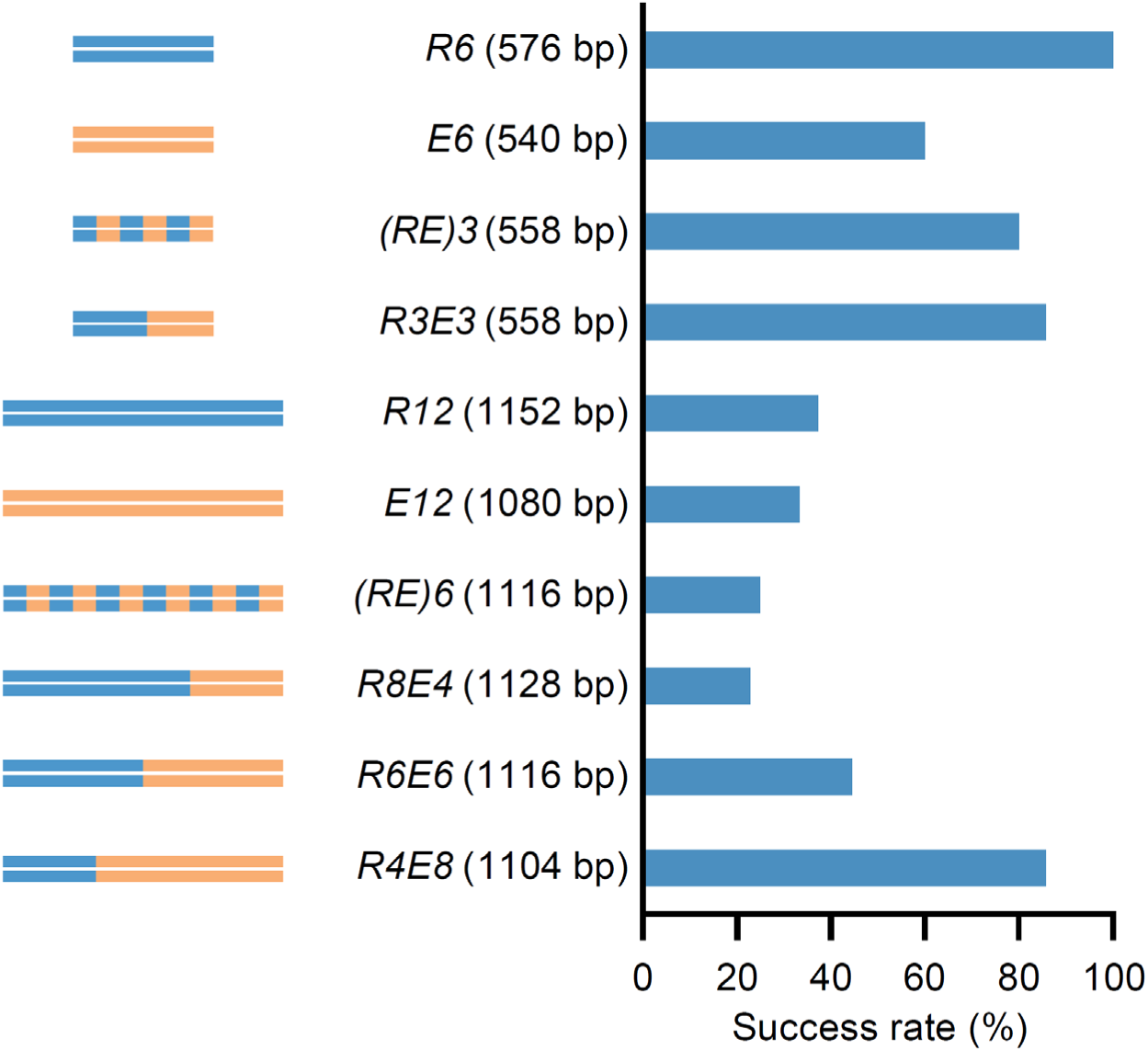
Success rates of various repetitive-sequence gene syntheses The success rates of gene syntheses were calculated from the positivity ratios obtained via colony PCR and Sanger DNA sequencing.

### 2.3. Characterization of the polymers prepared by SCRCA

Copolymerization or block copolymerization using two repeat units with different functions is a powerful method to develop multifunctional protein polymers;[25,26,42–44] however, fundamental studies are essential to determine the optimal compositions of such polymers. The conventional methods require at least two cloning operations per prototype copolymer[26,45]. In contrast, as shown in Figure 1b and c, SCRCA can produce polymers of similar lengths with different amino acid compositions all in a single cloning operation. Therefore, to confirm that polymers constructed with SCRCA can be used to elucidate the relationship between function and sequence, highly purified polymers were prepared using His-tag affinity resin (**Figure S8**, the polymers corresponding to genes are written in Roman type). By evaluating the temperature response of the polymers, we found that the polymer with a 2:1 ratio of RLP to ELP sequences (R8E4) exhibited the phase separation phenomenon over a wide temperature range, reflecting the characteristics of RLPs and ELPs (Figure 4). Notably, copolymer (RE)6 and block copolymer R6E6 containing the same amino acid composition exhibited similar temperature responses. This suggests that composition, rather than sequence, is related to the polymer function in the case of copolymerization of RLP and ELP. Such information could be efficiently obtained using SCRCA to accelerate the rational design of new multifunctional protein polymers.

**Figure 4.**
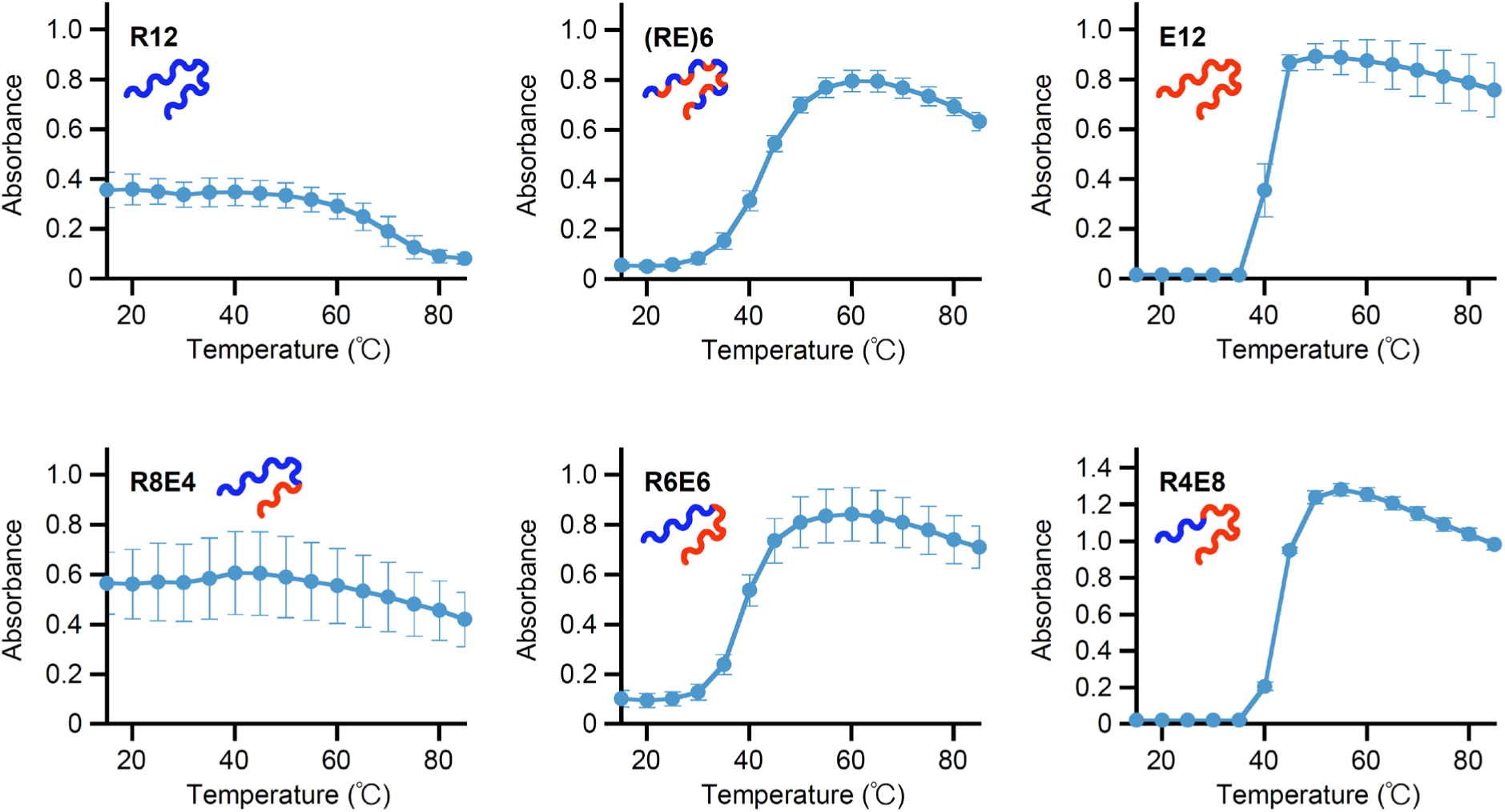
Temperature responses of the polymers prepared by seamless cloning of rolling-circle amplicons Phase separation behaviors of polymers R12, E12, (RE)6, R8E4, R6E6, and R4E8 with the increase in temperature. The samples were prepared in PBS (pH 7.4) containing 2 µM polypeptide and 8 mM residual urea. Each value represents the mean of three replicates, with error bars showing the standard errors.

### 2.4. Constructing of the Repeat Unit Library

Directed evolution is a promising technique to develop highly functional enzymes and antibodies[46], but to our knowledge, it has not been used to develop protein polymers. This may be due to the fact that mutagenesis methods suitable for low-complexity proteins such as protein polymers have not yet been explored. We considered that a repetitive-sequence gene library with repeat units containing mutations would be useful for directed evolution of protein polymers, and investigated an approach for the preparation of a ring-template mixture using ssDNA containing mixed bases (Figure 1d). ELP was selected as the model sequence; the <u>V</u> sites of <u>V</u>GVPG<u>V</u>GVPG<u>V</u>GVPG<u>V</u>GVPG<u>V</u>GVPG<u>V</u>GVPG can be replaced with a guest amino acid, which changes the temperature responsiveness of the ELPs[47] (Figure 5a). We designed an E-3NDT ssDNA oligo in which the three codons (X_1_, X_2_, and X_3_) encoding the <u>V</u> sites were replaced with NDT codons.[48] This codon encodes one of 12 diverse amino acids (Figure 5b).

**Figure 5.**
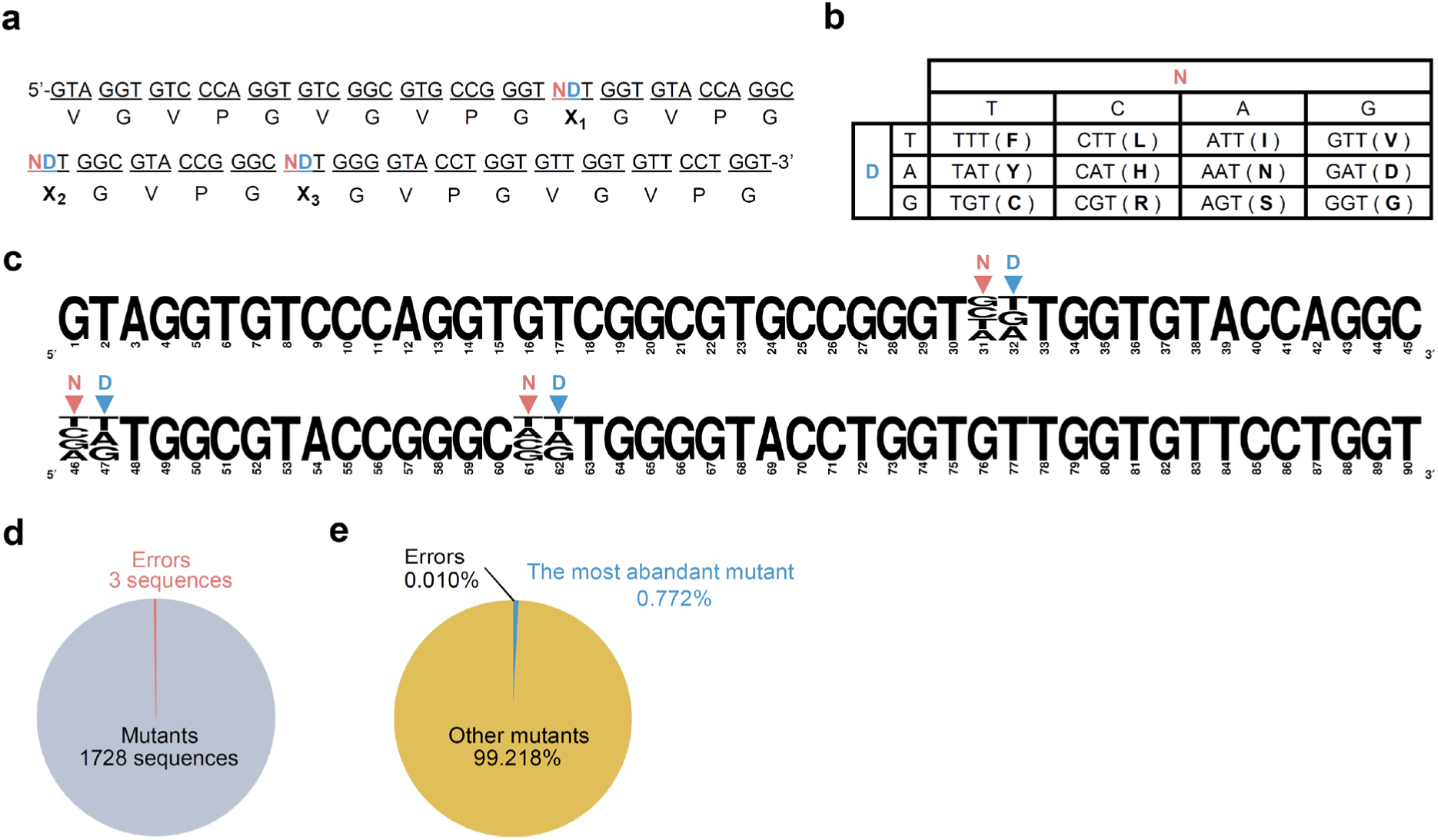
Construction of the repetitive-sequence gene library (a) Nucleotide sequence of the E-3NDT ring template library. (b) At each NDT codon site, one of 12 types of amino acids is encoded by the combination of bases. (c) Alignment result of the nucleotide repeat units found in the library constructed by RCA using the E-3NDT ring template library. The height of each nucleotide corresponds to its frequency. N and D represent the mixed base introduction position in E-3NDT ssDNA. (d) Ratios of mutants and errors among all sequences. Each value represents the mean of two replicates. The corresponding standard deviations are 0.7 and 1.4, respectively. (e) Abundance ratios of the most abundant mutant, other mutants, and error sequences. Each value represents the mean of two replicates. The corresponding standard deviations are 0.083%, 0.088%, and 0.005%, respectively.

To confirm the construction of the library, RCA products, which were presumed to have five repeats of the E-3NDT units, were analyzed using a next-generation sequencer. The nucleotide repeat units present at each end of the gene (the first at the 5′-end and the fifth at the 3′-end) could be read accurately. However, the sequence information at the second to fourth nucleotide repeat units was unreliable, probably owing to poor cluster formation due to the repetitive sequences. The counts of each sequence were almost the same in the first (at the 5′-end) and fifth (at the 3′-end) repeat units (R = 0.991), indicating that single nucleotide repeat units were repeated five times in most genes. Multiple alignments of the observed units confirmed that the E unit backbone was maintained (Figure 5c). In contrast, high complexity was observed at sites where mixed bases were introduced. While this library contained all the 1728 expected sequences (all combinations of the mixed bases), it contained few error sequences (Figure 5d). Notably, even the most abundant sequences represented only <0.9% of the total number of sequences, indicating that this library possessed excellent diversity (Figure 5e).

Genes having six repeats of the E-3NDT unit were introduced into the *E. coli* expression system mentioned above. As E-3NDT is based on the E unit, this can be termed an E6 mutant library. We examined 42 transformed colonies and selected 28 colonies with the correct gene lengths (**Figure S9**). Sanger sequencing revealed that most of the selected transformants carried E6 mutant genes with different amino acids (Figure 6a and b; the mutants are denoted by “m” and the colony number). Interestingly, block copolymer mutants with two or more different repeat units were obtained in addition to E6 mutants with the same unit repeated six times (as expected) (Figure 6b). We ascribe this to seamless cloning between the amplicons with ssDNA regions (Figure 6c). This phenomenon, which may also have occurred when constructing the genes comprising only one repeat unit, might facilitate the synthesis of long repetitive-sequence genes via SCRCA. Considering that polypeptide aggregation is affected by mean hydropathy and net charge,[49] the presence of block copolymer genes diversified the ELP library (Figure 6d).

**Figure 6.**
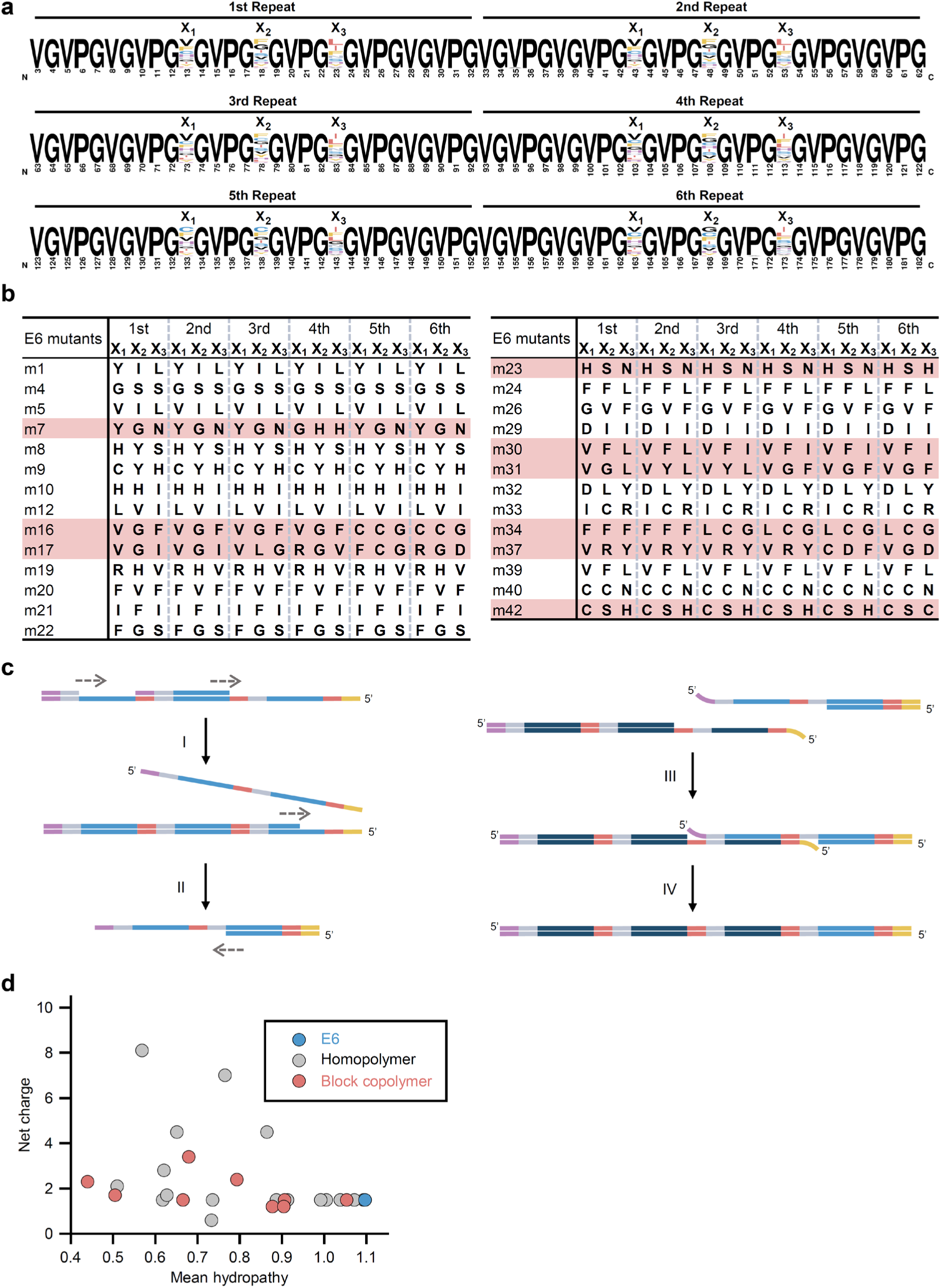
Composition of the E6 mutant library (a) Alignment result of the repetitive amino-acid-sequence parts (between 3-183 aa) of 27 E6 mutants fabricated by SCRCA using the E-3NDT ring template. The height of each amino acid corresponds to its frequency. The X_1_, X_2_, and X_3_ positions represent NDT codons in the E-3NDT ring template. (b) For 27 E6 mutants, the amino acids at X_1_, X_2_, and X_3_ are summarized. The mutants highlighted in red have a block copolymer sequence consisting of two or more repeat units. (c) Principle of block copolymer formation during the library construction. During isothermal amplification (described in Figure 2a), dsDNAs (I) with an ssDNA region at the 3′ end and (II) with an ssDNA region at the 5′ end are generated, respectively (blue, gray, and red lines: repetitive sequences; purple and yellow lines: overlapping sequences). (III) They anneal, and (IV) the nick is repaired during the In-Fusion reaction to generate a block copolymer gene. The dark blue line indicates a repeat unit that is partly different from that indicated by the blue line. (d) Mean hydropathy and net charges are plotted for wild type E6 (blue), 18 mutants with a homopolymer gene (gray), and 9 mutants with a block copolymer gene (red).

Recently, a library construction technique applicable to low-complexity sequences such as ELP was been reported, however, this technique presents challenges, such as transformation efficiency of less than 25% and low library diversity (the occupancy of the most abundant sequences is more than 30%)[50]. In contrast, libraries constructed using SCRCA have a transformation efficiency of approximately 70% and excellent diversity. In addition, the cost associated with SCRCA is only one-tenth that of the previously reported method because the introduction of mixed bases is free. These advantages greatly reduce the hurdles in applying evolutionary molecular engineering to the development of functional protein polymers.

### 2.5. Development of Desired Functional Protein Polymer by Directed Evolution

Finally, we tested whether the repeat unit library could be used to rapidly develop the desired protein polymers. As a development target, we selected a multi-response ELP that is soluble at low temperatures (4 °C, pH 7.4), insoluble near body temperature (37 °C, pH 7.4), and soluble in the environment around cancer cells (37 °C, pH 6.5). Such protein polymers are useful vehicles for drug delivery to cancer cells.[24,51] To develop polymers with these complex properties, molecules with LCSTs in the range of 4–37 °C at pH 7.4 and >37 °C at pH 6.5 must be designed (Figure 7a). The rational design of such polymers from the basic ELP sequence requires several years, as factors such as amino acid substitution, chain length, and influence of the operating environment require careful examination to determine their effect on polymer responsiveness to temperature and pH changes.[52–55]

**Figure 7.**
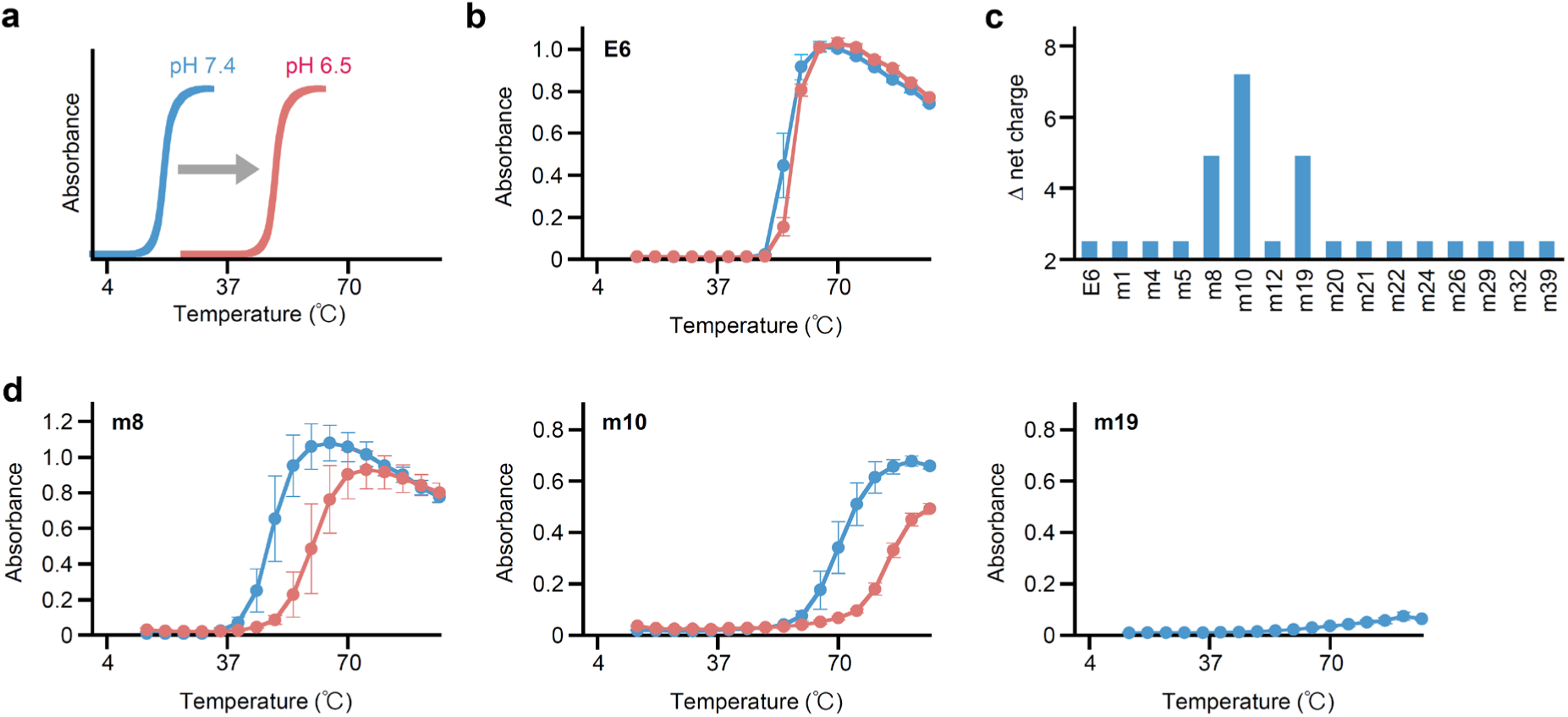
First screening of the directed evolution of the elastin-like polymer (a) The development target must have a transition temperature between 4 °C and 37 °C in a biological environment (pH 7.4) and >37 °C in a slightly acidic environment (pH 6.5). (b) Temperature and pH responses of E6. The sample was prepared in phosphate-buffered saline (PBS) solutions containing 5 μM E6 (blue line: pH 7.4; red line: pH 6.5). Each value represents the mean of three replicates, with error bars showing the standard errors. (c) Net charges at pH 7.4 and pH 6.5 calculated based on the respective amino acid sequences. (d) Temperature and pH responses of the three selected E6 mutants, i.e., m8, m10, and m19. The samples were prepared in PBS solutions containing 5 μM polymer (blue line: pH 7.4, red line: pH 6.5). Each value represents the mean of three replicates, with error bars showing the standard errors. The temperature response of m19 at pH 6.5 was not evaluated because this mutant did not show a lower critical solution temperature below 95 °C.

In the directed evolution experiment, we aimed to develop the desired polymer by optimizing the <u>V</u> positions of E6, which has an LCST higher than body temperature and does not respond to pH changes (Figure 7b). A problem encountered during screening is that the temperature responsiveness of ELP is easily affected by its concentration and the presence of contaminants.[52] Therefore, prior to screening, we predicted the polymer functions from the amino acid sequences and then selected some mutants to reduce the number of samples. By evaluating the temperature responsiveness of the selected mutants after purification, the improvement in the function can be compared.

The screening of the first directed evolution round involved searching for protein polymers that exhibited temperature and pH responsiveness in the E6 mutant library. Cysteine-containing mutants (m9, m33, and m40) were excluded owing to their potential to form irreversible aggregates. Block copolymers were also excluded because of the complexity of the next-generation library design. Consequently, three mutants, m8, m10, and m19 whose net charge changed significantly in response to a small change in pH, were selected and purified (Figure 7c **and S10**). The mutants m8 and m10 exhibited both temperature and pH responsiveness (Figure 7d). However, they were water-soluble near body temperature.

The first-round results (Figures 6b**, 7c, and 7d**) reveal that substitution of histidine results in the response to the small pH change. Lowering LCST while maintaining pH responsiveness would require the introduction of functional groups that interact with histidine, such as tyrosine, phenylalanine, and histidine[56]. Therefore, we designed the m10-derived sequence m10-3YWY, which has YWY codons encoding one phenylalanine, tyrosine, leucine, and histidine in the remaining three <u>V</u> sites (positions X_4_, X_5_, and X_6_) (Figure 8a and b).

**Figure 8.**
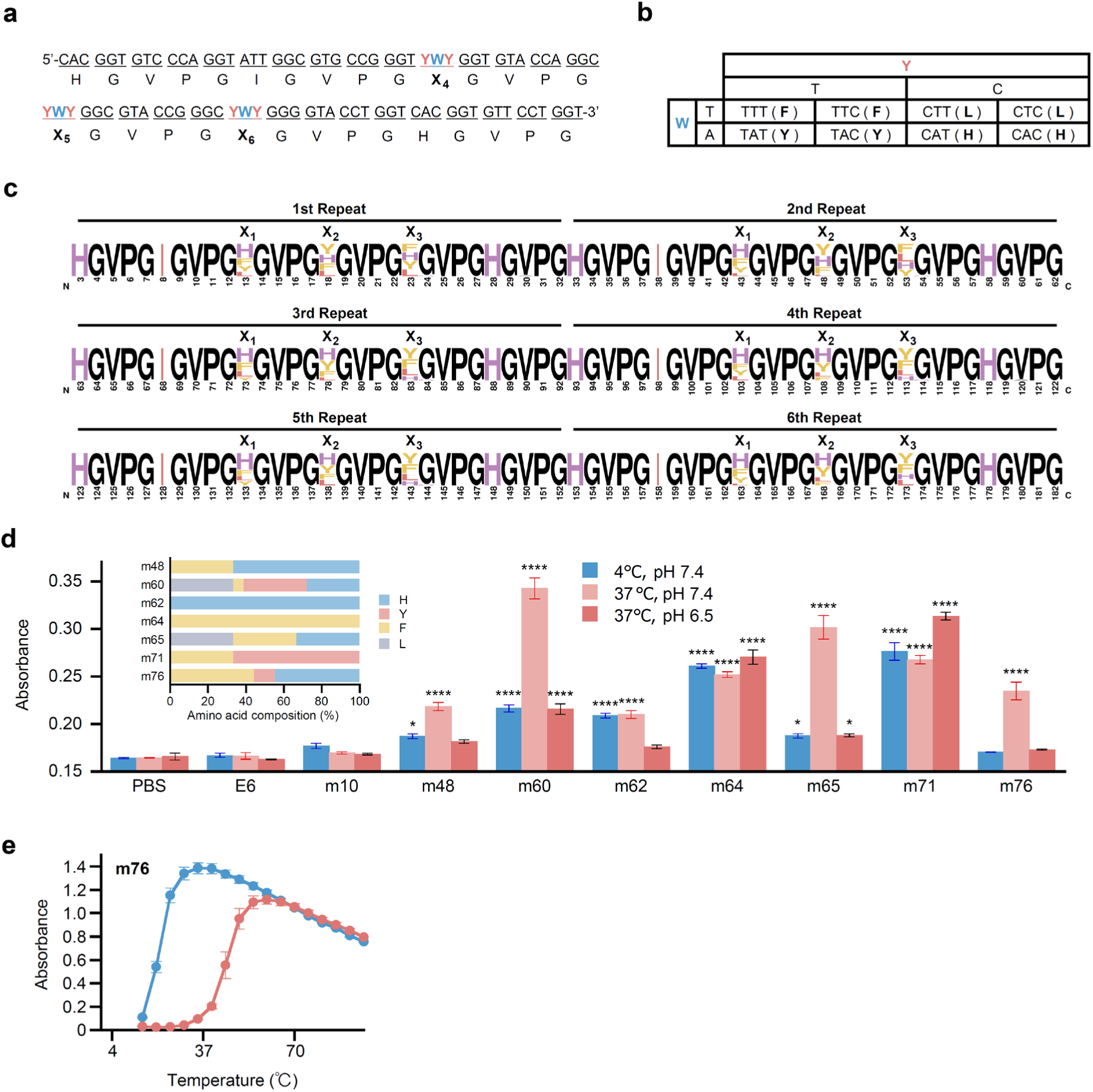
Second screening of the directed evolution (a) Nucleotide sequence of the m10-3YWY ring template. (b) At each YWY codon site, one of four types of amino acids is encoded by the combination of bases. (c) Alignment result of repetitive amino-acid-sequence parts (in the range of 3 to 183 aa) of 27 m10 mutants prepared by SCRCA using the m10-3YWY ring template. The height of each amino acid corresponds to its frequency. The X_4_, X_5_, and X_6_ positions represent YWY codons in the m10-3YWY ring template. (d) Results of simple turbidity measurements. Seven m10 mutants, i.e., m48, m60, m62, m64, m65, m71, and m76, were selected based on their unique amino acid compositions (insert). The samples were prepared in PBS (pH 7.4) containing 5 μM polymer and < 270 mM residual urea. Turbidity was measured at 4 °C and 37 °C, and then, the pH was adjusted to 6.5 by adding lactic acid. Each value represents the mean of three replicates, with error bars showing the standard errors. Statistical differences were determined using one-way ANOVA with Dunnett’s multiple comparisons post-test. Differences in turbidity (at pH 7.4 and 4 °C) relative to the PBS blank: no asterisk *P* > 0.1; * *P* < 0.1; **** *P* < 0.0001. (e) Temperature and pH responses of m76. The samples were prepared in PBS solutions containing 5 μM polymer and 80 mM residual urea (blue line: pH 7.4, red line: pH 6.5).

Among the 37 transformants examined using colony PCR and Sanger sequencing (**Figure S11**), 27 mutants possessed the m10 backbone with different amino acids at positions X_4_, X_5_, and X_6_ (Figure 8c **and S12**). This indicates that a template design can be used to construct a protein polymer library that inherits the characteristics of the useful sequences identified in the previous round.

To reduce the library size, we selected seven unique mutants based on their amino acid composition (Figure 8d) and purified them (**Figure S13**). We measured turbidity in low-temperature (4 °C, pH 7.4), biological (37 °C, pH 7.4), and slightly acidic (37 °C, pH 6.5) environments on a microplate scale. Based on the result (Figure 8d), we selected m76, which agglutinated only at 37 °C, pH 7.4, as a strong candidate; its temperature and pH responses were close to the target properties (Figure 8e). This result indicates that the SCRCA-assisted directed evolution strategy could allow for the development of a desired protein polymer with a few transformation operations and within several months.

## 3. Conclusion

We developed a repetitive-gene synthesis method that combines RCA and seamless cloning and demonstrated that various genes encoding protein polymers can be easily synthesized. The SCRCA method has a high success rate of gene synthesis and excellent workability, and block copolymer genes, which normally require multiple cloning steps, can be constructed in a single cloning operation. As the operation days and cost of SCRCA are comparable to those of standard gene synthesis, SCRCA is expected to be a fundamental technology that will accelerate the rational design of functional protein polymers.

We also demonstrated that protein polymer libraries with different repetitive units can be easily constructed by introducing mixed bases into the template ssDNA. The library construction using SCRCA achieved superior transformation efficiency and diversity in relation to that using the state-of-the-art method. Furthermore, by combining this library construction technique with the directed evolution concept, we proved that the development of a highly functional protein polymer can be easily achieved without the need for diligent research. This development strategy may be suitable for further functionalization of conventional protein polymers and for the search for polymers with unknown functions. Considering its suitability for the directed evolution of low-complexity sequences, this strategy may also be used to understand diseases involving intrinsically disordered proteins[57,58] and to develop bioproduction processes using intracellular phase separation.[59] Future research will focus on efficiently screening desired protein materials and transformants. As proteins and polypeptides have a wide range of applications, a screening method that is suitable for development purposes must be proposed. Further studies will advance SCRCA to a technology that contributes substantially to multiple fields.

## 4. Methods

### 4.1. Synthesis of the ssDNA Ring

DNA was synthesized by Eurofins Genomics, Inc. (Tokyo, Japan). The ssDNA ring template was designed based on the repeat-unit sequence and synthesized as 5′-phosphorylated ssDNA. The forward and reverse primers were designed based on the beginning and end sequences of the repeat unit, respectively. By designing the reverse primer to anneal to both ends of the 5′-phosphorylated ssDNA, the reverse primer was also used as a split oligo for ssDNA cyclization. The combinations of the ssDNA and primers used in this study are summarized in Table S1.

A mixture of 2 µL 50 µM 5′-phosphorylated ssDNA, 4 µL 50 µM reverse primer, and 29 µL milli-Q water was heated at 95 °C for 2 min. After cooling at 4 °C, 1 µL T4 DNA ligase (400,000 cohesive end units/mL, New England Biolabs) and 4 µL 10× T4 DNA ligase buffer were added, followed by incubation at 20 °C overnight. The product was purified using a GenElute PCR Clean-up Kit (Sigma-Aldrich, St Louis, MO, USA). Purity and concentration were determined using a Nanodrop 2000 device (Thermo Fisher Scientific, Waltham, MA, USA).

To prepare the RE ring template, a split DNA (ACCAGGAACACCATCACCACGACC) was designed based on both ends of the 5′-phosphorylated ssDNAs of the R and E rings. A mixture of 2 µL of each of the 50 µM 5′-phosphorylated ssDNAs, 4 µL of the 50 µM split DNA, 4 µL of the 50 µM reverse primer, and 23 µL milli-Q water was prepared. Cyclization was performed as described above.

### 4.2. RCA

A reaction solution of 100 µL containing 0.32 units/µL Bst DNA polymerase large fragment (New England Biolabs), 0.5 µM forward primer, 0.5 µM reverse primer, 0.2 ng/µL ssDNA ring template, 1.5 mM dNTP mix, 1× Thermopol buffer, and 2 mM magnesium sulfate was prepared. Isothermal amplification was performed at 60 °C for 12 h. A 1.5% TBE agarose gel was used to confirm the amplification products. The agarose gels were stained with ethidium bromide after electrophoresis and photographed using the Dolphin-Doc imaging system (Wealtec Corp., Sparks, NE, USA). Band positions were calculated using CS Analyzer 3.0 (ATTO Corporation, Tokyo, Japan).

### 4.3. Cloning and Transformation

For size selection of the DNA fragments, 2.5% TBE agarose gels containing 0.01% (v/v) SYBR Safe DNA were used. After electrophoresis, the band positions were checked using a blue-light illuminator, and the band corresponding to the desired length was cut using a cutter. The DNA fragments were purified using the FastGene Gel/PCR Extraction Kit (Nippon Genetics, Tokyo, Japan). The purity and concentration were determined using the Nanodrop 2000 device.

Expression vectors were constructed using an In-Fusion cloning kit (Takara Bio Inc., Shiga, Japan) according to the manufacturer’s instructions. A linear vector was constructed via inverse PCR using the pET22b vector and two primers (TGGCCGACTCATCATCACCACCACCAC and TTCATATGTATATCTCCTTCTTAAAGTTAAAC). The sequences of these primers were designed to add the amino acid sequences MK to the N-terminus and WPTHHHHHH to the C-terminus of protein polymers. *E. coli* BLR (DE3)-competent cells prepared in the laboratory were used for the transformation.

### 4.4. Colony PCR, Sanger DNA Sequencing and Mass Spectrometry

The T7-promoter primer (TAATACGACTCACTATAGG) and T7-terminator primer (GCTAGTTATTGCTCAGCGG) were used for colony PCR and Sanger DNA sequencing. Colony PCR was performed according to the instructions for the Go Taq Green Mix (Promega, Madison, WI, USA) except that the annealing temperature was set to 50 °C, and the elongation time was set to 2 min for short genes (approximately 550 bp) and 3 min for long genes (approximately 1100 bp).

Sanger DNA sequencing was performed by Fasmac Co. Ltd (Atsugi, Japan). BLAST (https://blast.ncbi.nlm.nih.gov/Blast.cgi) was used to analyze the sequence results. For genes with repetitive sequences of >1000 bp, the results of decoding in the 5′→3′ and 3′→5′ directions were combined (duplicated portions were conformed to be >60 bp). The positivity rate (%) was calculated as [number of positive results from colony PCR / number of tests for colony PCR] × [number of positive results from Sanger DNA sequencing / number of tests for DNA sequencing].

For the protein polymers whose repeat counts could not be confirmed by Sanger sequencing alone [R12, E12, and (RE)6], masses were measured via matrix-assisted laser desorption ionization time-of-flight mass spectrometry (MALDI-TOF-MS). The measurements were performed by Apro Science (Tokushima, Japan).

### 4.5. Preparation of Block Copolymer Genes

For simultaneous seamless cloning, RLP-block genes were prepared via RCA with a forward primer (GATATACATATGAAAGGGCGCGGTGACTCTCC) and a reverse primer (GACACCTACTGAGTAAGGTGAATCACCACGACC). These genes have an overlapping sequence at their 3′-end that recognizes the 5′-end of ELP-block genes. The ELP-block genes were prepared using RCA with a forward primer (TACTCAGTAGGTGTCCCAGGTGTCGG) and a reverse primer (ATGATGAGTCGGCCAACCAGGAACACCAACACCAGGTAC). The desired band of the block gene was cut out from the gel and purified. Two block genes and a linear vector were linked via an In-Fusion reaction.[41]

### 4.6. Expression and Purification

Transformants were inoculated into a 200 mL ZYP-5052 medium and incubated at 25 °C for 2–3 d. After harvesting by centrifugation (9000 *×g*, 5 min), the bacteria were stored at −30 °C. Protein polymers R12, E12, (RE6), R8E4, R6E6, and R4E8 were purified using His-tag affinity resin (His GraviTrap or Ni Sepharose 6Fast Flow, GE Healthcare, Chicago, IL, USA). Initially, 10 mL binding buffer (20 mM phosphate buffer (pH 7.4) containing 4 M urea, 0.5 M NaCl, and 40 mM imidazole) was added to the defrosted bacteria, and the bacteria were then sonicated on ice. After insoluble matter was removed via centrifugation, the supernatant was incubated at 37 °C for 10 min. Centrifugation was again performed to remove the generated insoluble matter, and the supernatant was passed through a 0.2 µm filter for sterilization. Next, affinity purification was performed according to the manufacturer’s instructions, except that 0.04 and 4 M urea were added to the binding and elution buffers, respectively. The eluate was concentrated using an Amicon Ultra 10 kDa column (Millipore, Burlington, MA, USA). Protein polymers E6, m8, m10, and m19 were purified via the inverse transition cycling (ITC) method.[15] The purified polypeptides were dissolved in the milli-Q water or phosphate-buffered saline (PBS). Protein polymers m48, m60, m62, m64, m65, m71, and m76 were purified using the His-tag affinity resin and then further purified using the ITC method. The purified products were dissolved in 20 mM phosphate buffer (pH 7.4) containing 4 M urea to maintain their high concentrations and stored at −20 °C.

### 4.7. Verification of Protein Polymer Purity

Purified protein polymer concentrations were calculated by measuring absorbance at 280 nm. The molar absorption coefficient was determined using the method reported by Pace et al.[60] To check for contamination by other proteins, sodium dodecyl sulfate-polyacrylamide gel electrophoresis (SDS-PAGE) was performed, and the gels were stained with Coomassie brilliant blue.

### 4.8. Evaluation of Temperature Response

Absorbance was measured at 350 nm using a V-650 spectrophotometer (Jasco, Tokyo, Japan) connected to a temperature controller. PBS at pH 7.4 was used as the blank. To facilitate the adjustment of the polymer concentration, the measurement solution was prepared by diluting a concentrated solution. Some concentrated solutions contained 4 M urea to prevent phase separation at room temperature. The residual urea concentration in the measurement solution is presented in the figure legends. The temperature was increased by 1 °C per min. PBS at pH 6.5 was prepared by adding 22 μL of 50-fold diluted lactic acid to 1000 μL of the measurement solution.

### 4.9. Next-generation Sequencing Analysis

MiSeq Illumina sequencing (Illumina, San Diego, CA, USA) and data analysis were performed by Hokkaido System Science Co., Ltd (Hokkaido, Japan). The sequencing samples were synthesized via RCA using a set of primers (TCGTCGGCAGCGTCAGATGTGTATAAGAGACAGTGTAGGTGTCCCAGGTGTCGG and GTCTCGTGGGCTCGGAGATGTGTATAAGAGACAGACCAGGAACACCAACACCAGGTAC) with linker sequences at their 5′-ends. For size selection, agarose gel electrophoresis was conducted, and fragments corresponding to five repeats were excised from the gel. After purification, the samples were submitted to the company. Analysis revealed that the middle portions of the gene sequences were less reliable; therefore, only the sequences that were 90 bp from the 5′-ends and 90 bp from the 3′-ends were included in the analysis. However, as sequences with low counts lack reliability, those with six or more counts were targeted for analysis. All sequences in a library were aligned in parallel using Weblogo (weblogo.berkeley.edu/logo.cgi).

### 4.10. Mean Hydropathy and Net Charge

Mean hydropathy and net charge were calculated using Expasy (https://web.expasy.org/protscale/) and PROTEIN CALCULATOR 3.4 (https://protcalc.sourceforge.net/), respectively. Calculations were performed using default settings.

### 4.11. Simple Turbidity Test

Polymer solutions of 150 µL (5 µM polypeptide in PBS, pH 7.4) were prepared in the cells of 96-well microplates (clear bottom, half area). As the measurement solutions were prepared by diluting the polymer concentrates with 4 M urea, the solutions contained 40–270 mM of urea, which should have little effect on the transition temperature of ELPs.[61] The solutions were incubated at 4 °C for 30 min, and absorbance was measured at 350 nm. The solutions were further incubated at 37 °C for 30 min, and absorbance was measured again. Then, 3.3 µL of 50-fold diluted lactic acid was added to reduce the pH to 6.5. The solution was again incubated at 37 °C for 30 min, and absorbance was measured at 350 nm. PBS was used as a blank. The experiments were performed on three cells under identical conditions.

### 4.12. Statistical Analysis

The correlation coefficients were calculated and statistical tests were performed using KaleidaGraph Version 4.5.3 (Synergy Software, Eden Prairie, MN, USA). One-way ANOVA with Dunnett’s test was used for multiple comparisons. Differences with *P* < 0.1 were considered statistically significant for improving screening accuracy.

## Supporting Information

Supporting Information is available from the Wiley Online Library or from the author.

## Acknowledgments

This study was partly supported by a Grant-in-Aid for Early-Career Scientists (No. 22K14549) and Grant-in-Aid for Transformative Research Areas (No. 24H01378) from the Japan Society for the Promotion Science (JSPS), a project (No. JPNP20004) subsidized by the New Energy and Industrial Technology Development Organization (NEDO), the project “Research project for sericultural bio-industry” (No. JP22680575) commissioned by the Ministry of Agriculture, Forestry and Fisheries (MAFF), and the Research Collaboration Program (No. AM2100) of the National Institute for Materials Science (NIMS).

## Data availability statement

The data sets in this study are available within the source data.

