## Supporting information for "A Novel Gene Synthesis Platform for Designing Functional Protein Polymers"

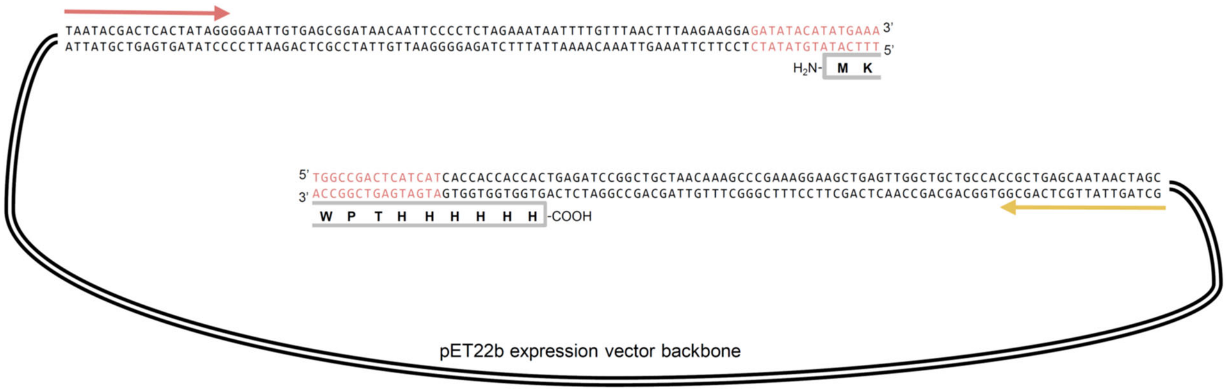


**Figure S1. Linear vector for protein polymer expression**

Red letters indicate overlapping sequences. Sequences at both ends of the linear vector were designed to add MK to the N-terminus and WPTHHHHHH to the C-terminus of the expressed protein polymers. Red and yellow arrows indicate primer annealing positions for colony PCR and Sanger DNA sequencing.

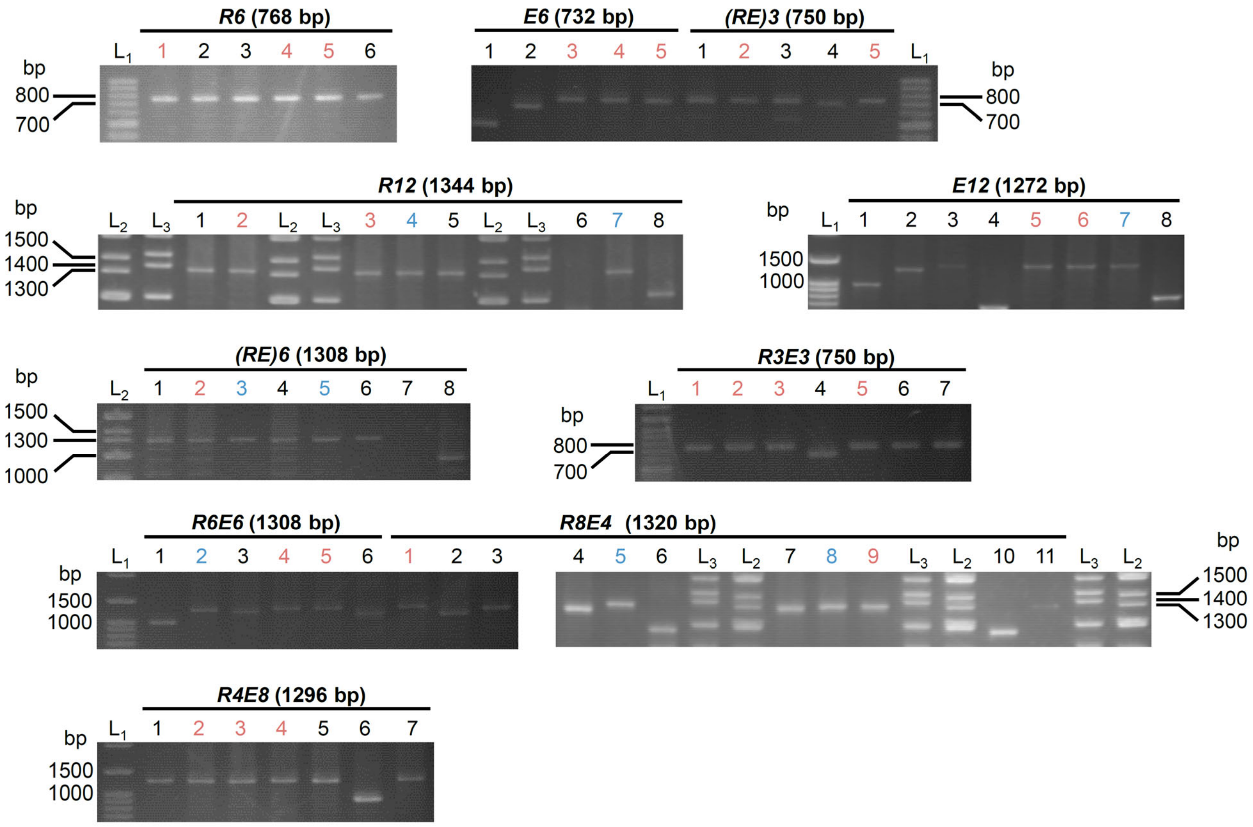


**Figure S2. Colony PCR results of transformants with various repetitive-sequence genes**

Transformants presumed to contain the gene of interest were selected using colony PCR. The size of the colony PCR products was confirmed by electrophoresis using a 1.5% agarose gel. [$L_1$: 100 bp DNA ladder Plus (Nippon Genetics); $L_2$: Gene ladder Wide 1 (Nippon Gene); $L_3$: Wide-Range DNA Ladder (Takara Bio, Shiga, Japan)]. The number of bases in parentheses represents the theoretical length of the colony PCR product. For the transformants identified by red and blue font, the gene sequences were confirmed by Sanger DNA sequencing, which revealed that the red-font transformants had a correct repetitive sequence.

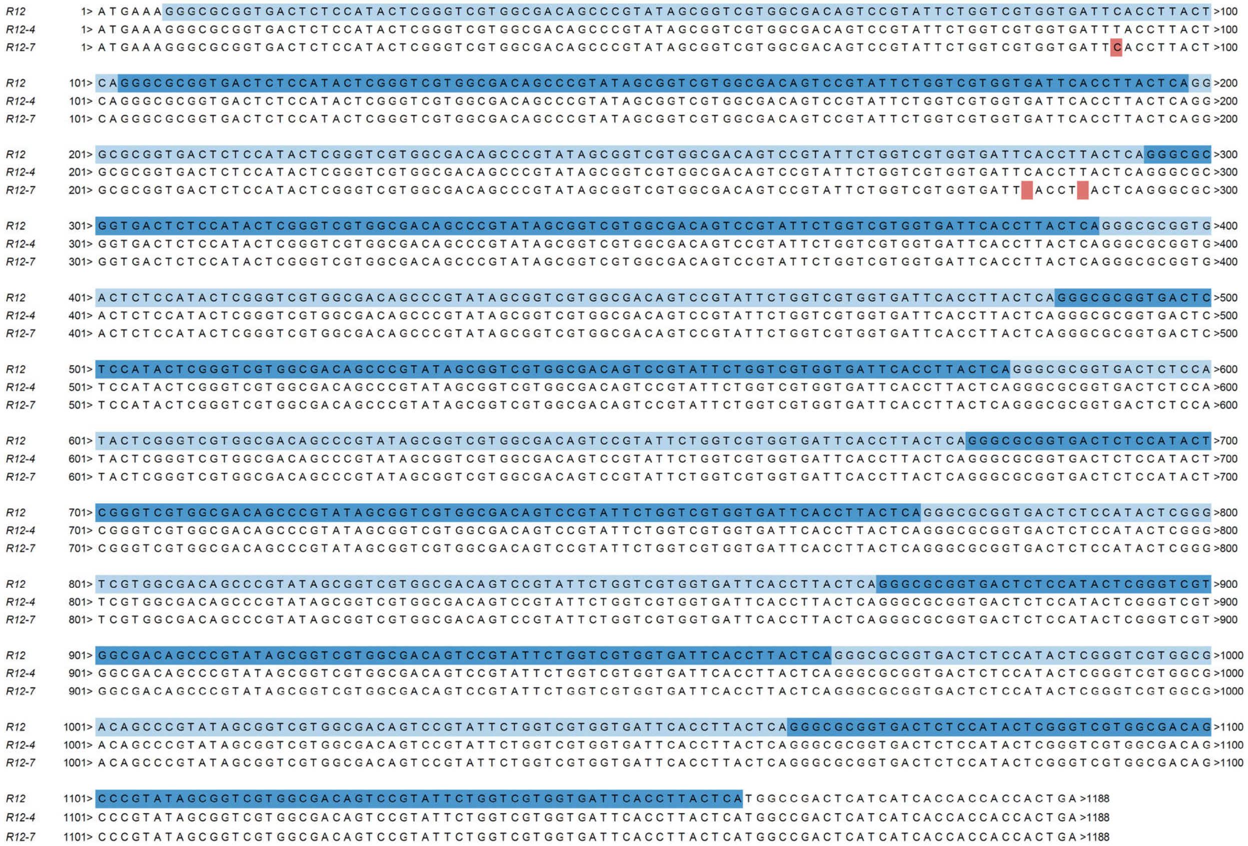

**Figure S3. Alignment of the correct open reading frame and the error nucleotide sequences of *R12***

Nucleotides marked in red differ from those present in the designed sequence. To easily identify repetitive sequences, odd repetitions are indicated in light blue and even repetitions in blue.

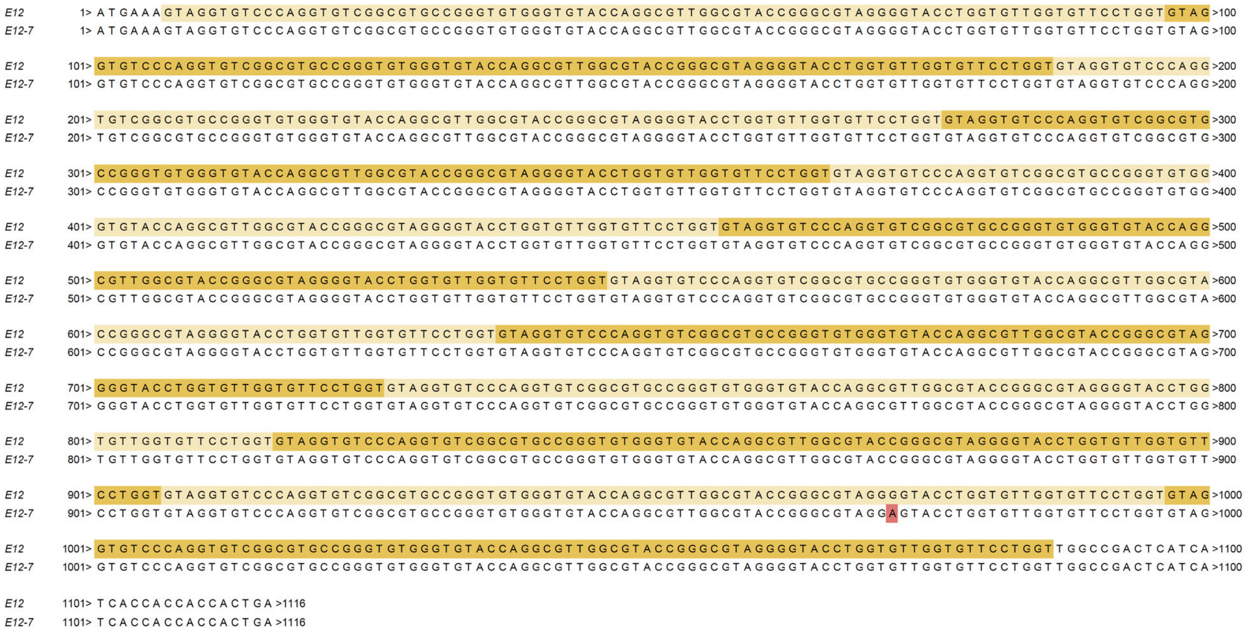

**Figure S4. Alignment of the correct open reading frame and the error nucleotide sequences of *E12***

Nucleotides marked in red differ from those present in the designed sequence. To easily identify repetitive sequences, odd repetitions are indicated in light yellow and even repetitions in yellow.

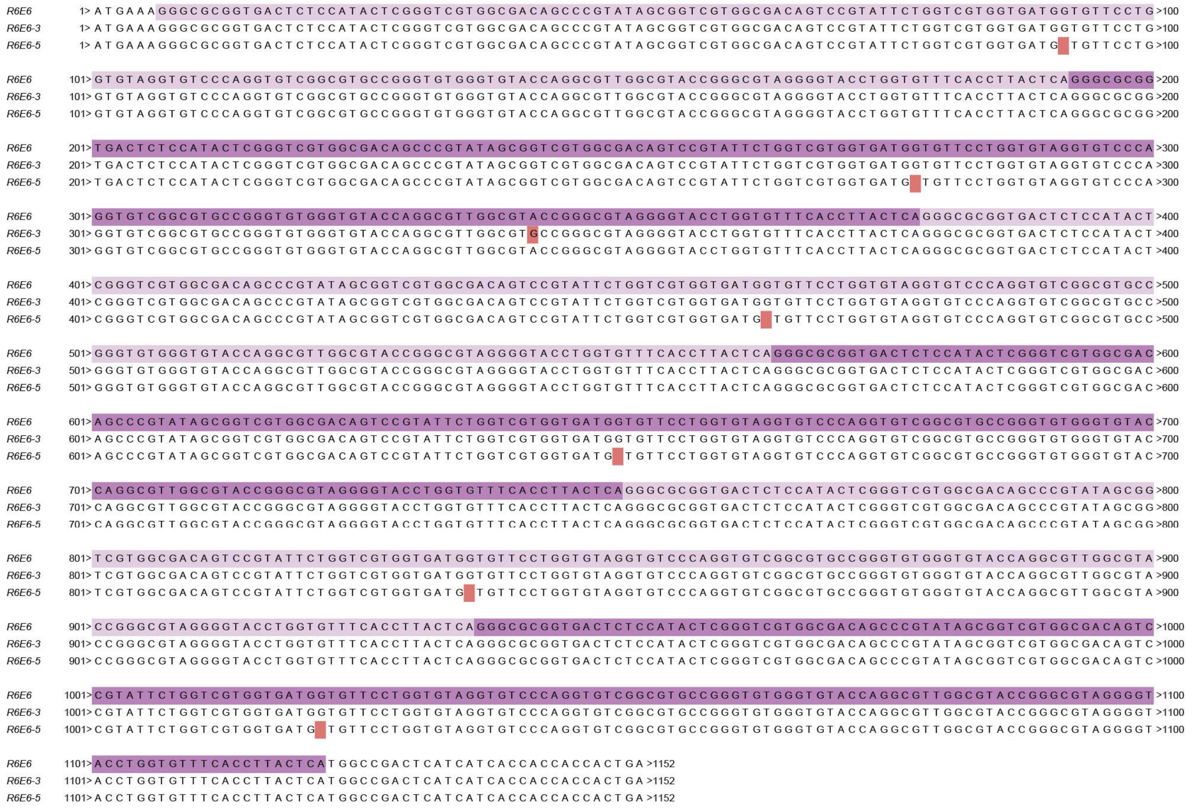

**Figure S5. Alignment of the correct open reading frame and the error nucleotide sequences of *(RE)6***

Nucleotides marked in red differ from those present in the designed sequence. To easily identify repetitive sequences, odd repetitions are indicated in light purple and even repetitions in purple. The gaps seen in *(RE)6*-5 can be attributed to an incomplete template.

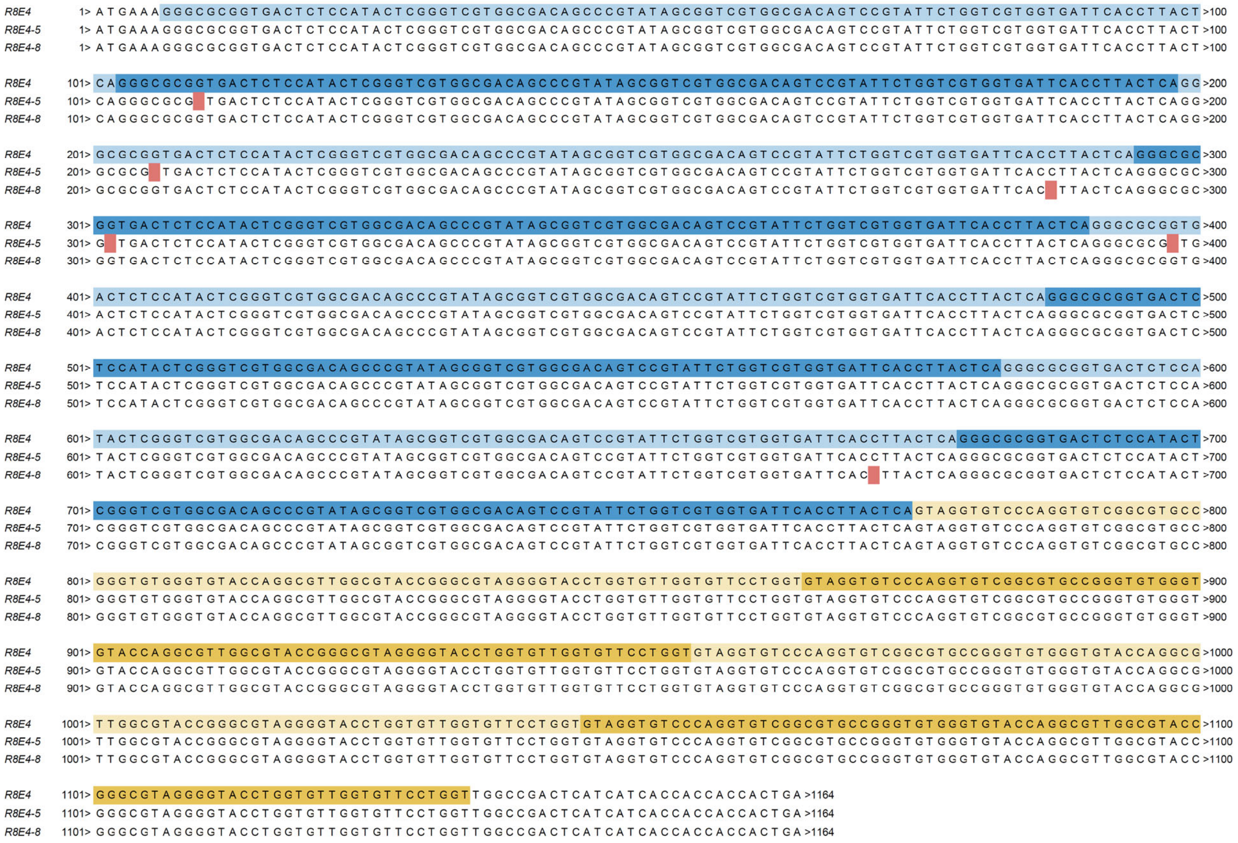

**Figure S6. Alignment of the correct open reading frame and the error nucleotide sequences of *R8E4***

Nucleotides marked in red differ from those present in the designed sequence. To easily identify repetitive sequences, odd repetitions of the R repeats are indicated in light blue and the even repetitions in blue, and odd repetitions of the E repeats are indicated in light yellow and the even repetitions in yellow. The gaps seen in *R8E4*-5 and *R8E4-8* can be attributed to an incomplete template.

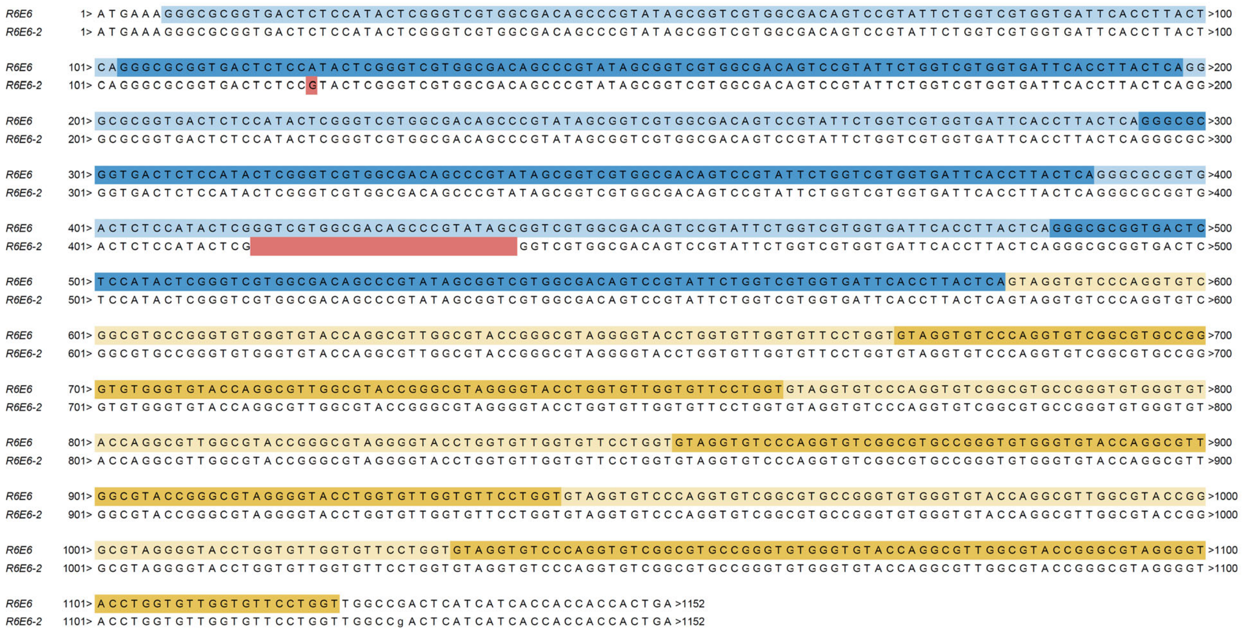

**Figure S7. Alignment of the correct open reading frame and the error nucleotide sequences of *R6E6***

Nucleotides marked in red differ from those present in the designed sequence. To easily identify repetitive sequences, odd repetitions of the R repeats are indicated in light blue and the even repetitions in blue, and odd repetitions of the E repeats are indicated in light yellow and the even repetitions in yellow. The continuous 24-base gap in *R6E6*-2 was assumed to be an error caused by secondary structure formation or cloning failure.

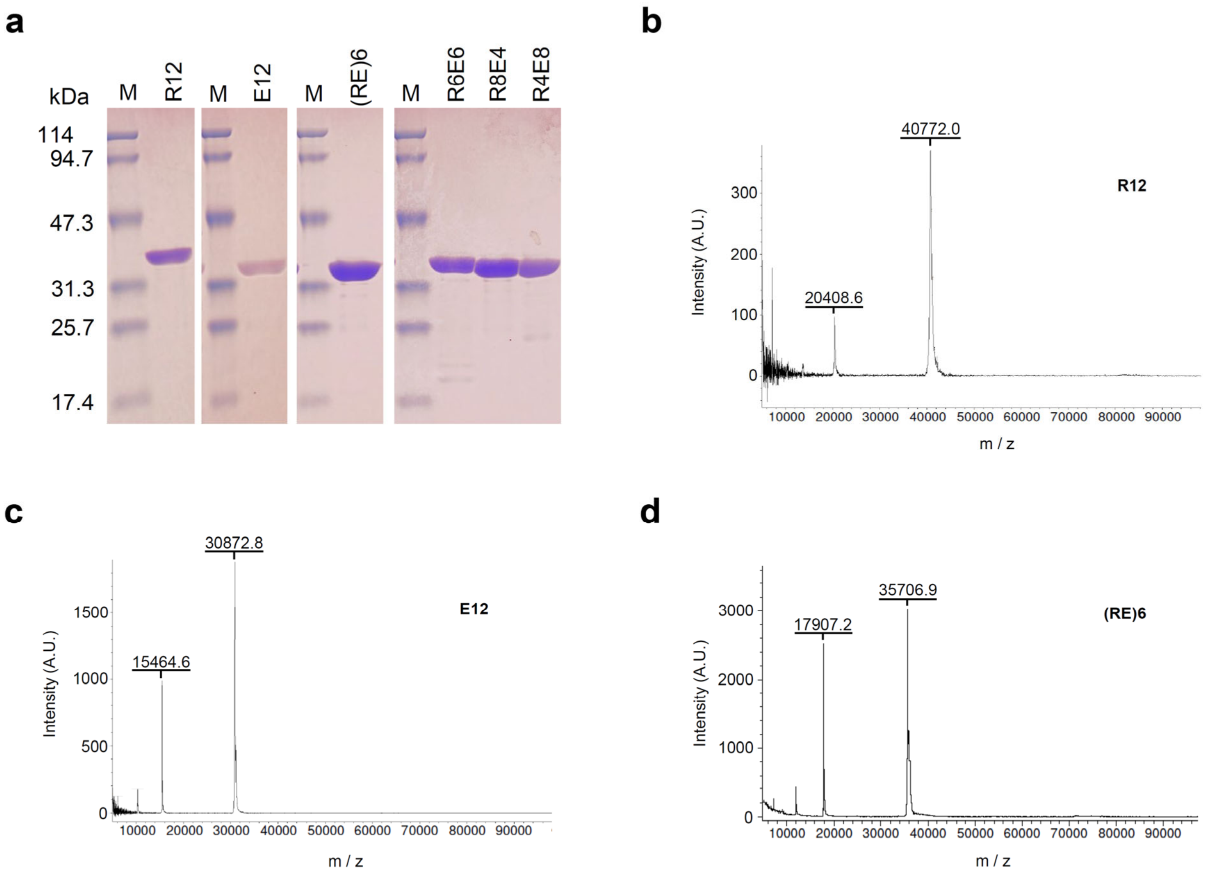


**Figure S8. Molecule sizes of the purified protein polymers**

(a) Lane M contains EzStandard PrestainBlue molecular-weight marker (ATTO, Tokyo, Japan). SDS-PAGE results confirmed that R12, E12, (RE)6, R8E4, R6E6, and R4E8 were highly purified. (b-d) Sanger DNA sequencing confirmed the repeat numbers of R8E4, R6E6, and R4E8, but could not confirm those of R12, E12, and (RE)6. Therefore, their masses were measured via MALDI-TOF-MS. The theoretical molecular weights of (b) R12, (c) E12, and (d) (RE)6 were 40817, 30949, and 35883, respectively; therefore, all three had masses within 0.5% of the theoretical values. This indicates that R12, E12, and (RE)6 were constructed as designed.

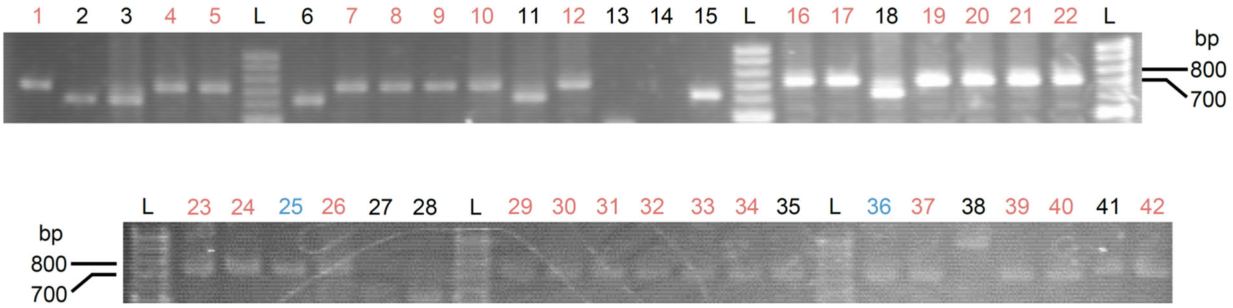


**Figure S9. Colony PCR results of the transformant library**

The transformants indicated using red and blue fonts were presumed to contain the target gene using colony PCR. The lengths of the colony PCR products were confirmed by electrophoresis using a 1.5% agarose gel (L: 100 bp DNA ladder Plus). The theoretical length is 732 bp. Sanger DNA sequencing revealed that the red transformants were E6 mutants. No analyzable sequence result was obtained from the No. 35 colony.

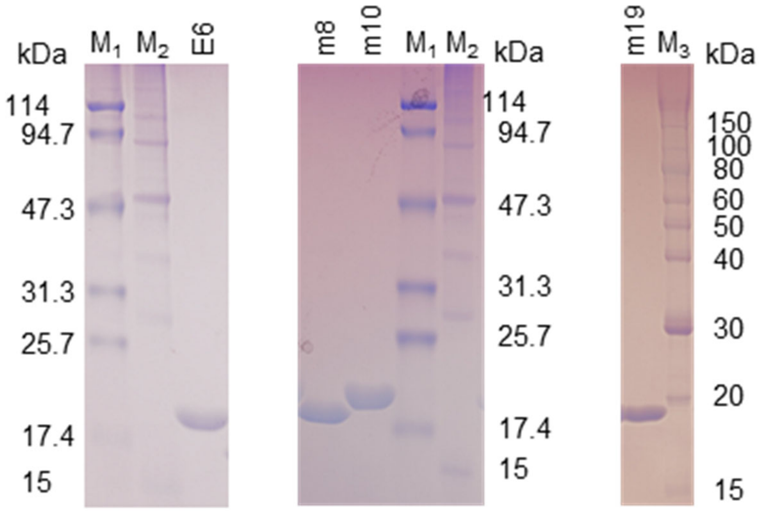


**Figure S10. SDS-PAGE images of the E6 mutants**

Lanes $M_1$, $M_2$, and $M_3$ contain EzStandard PrestainBlue molecular-weight marker (ATTO), Broad Range Protein Molecular Weight Markers (Promega, Madison, USA), and SIMASIMA Unstained Broad Range Protein Ladder (Cosmo Bio, Tokyo, Japan), respectively. No bands of contaminant proteins were observed, indicating that high-purity E6, m8, m10, and m19 were prepared.

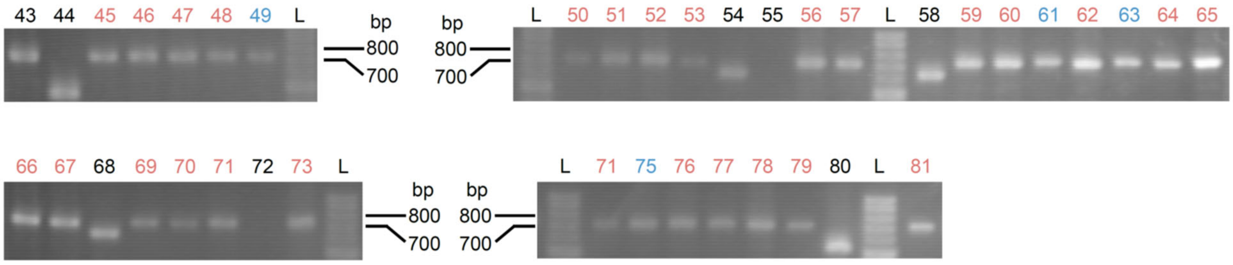


**Figure S11. Colony PCR results of the transformants obtained for the second round**

Transformants presumed to contain the target gene were selected using colony PCR. The length of the colony PCR products was confirmed by electrophoresis using a 1.5% agarose gel (L: 100 bp DNA ladder Plus). The theoretical length is 732 bp. The transformants indicated in red and blue fonts were determined to contain inserted DNA fragments of similar length as the target repeat sequence. DNA sequencing revealed that the red-font transformants were m10 mutants. No analyzable sequence result was obtained from the No. 43 colony.

| m10 mutants | 1st | | | 2nd | | | 3rd | | | 4th | | | 5th | | | 6th | | |
|---|---|---|---|---|---|---|---|---|---|---|---|---|---|---|---|---|---|---|
| | $X_4$ | $X_5$ | $X_6$ | $X_4$ | $X_5$ | $X_6$ | $X_4$ | $X_5$ | $X_6$ | $X_4$ | $X_5$ | $X_6$ | $X_4$ | $X_5$ | $X_6$ | $X_4$ | $X_5$ | $X_6$ |
| m45 | L | H | Y | L | H | Y | L | H | Y | L | H | Y | L | H | Y | L | H | Y |
| m46 | Y | Y | L | Y | Y | L | Y | H | F | Y | H | F | Y | H | F | Y | H | F |
| m47 | Y | Y | L | Y | Y | L | Y | Y | L | Y | Y | L | Y | Y | L | Y | Y | Y |
| m48 | H | H | F | H | H | F | H | H | F | H | H | F | H | H | F | H | H | F |
| m50 | H | Y | L | H | Y | L | H | Y | L | H | Y | L | H | Y | L | H | Y | L |
| m51 | H | L | H | H | L | H | H | L | H | H | L | H | H | L | H | H | L | H |
| m52 | L | H | H | L | H | H | L | L | Y | H | L | Y | H | L | Y | H | L | Y |
| m53 | L | L | L | H | F | F | H | F | F | H | F | F | H | L | L | H | L | L |
| m56 | L | H | L | L | H | L | L | H | L | L | H | L | L | H | L | L | H | H |
| m57 | H | Y | H | H | Y | H | H | Y | H | H | Y | H | H | Y | H | H | Y | H |
| m59 | Y | L | Y | F | Y | L | F | Y | L | Y | L | Y | Y | L | Y | F | Y | L |
| m60 | H | L | Y | H | L | Y | H | L | Y | H | L | Y | H | L | Y | F | L | Y |
| m62 | H | H | H | H | H | H | H | H | H | H | H | H | H | H | H | H | H | H |
| m64 | F | F | F | F | F | F | F | F | F | F | F | F | F | F | F | F | F | F |
| m65 | F | H | L | F | H | L | F | H | L | F | H | L | F | H | L | F | H | L |
| m66 | H | F | Y | H | F | Y | L | F | Y | L | F | Y | L | F | Y | H | Y | Y |
| m67 | H | Y | Y | H | Y | Y | H | Y | Y | H | Y | Y | H | Y | Y | H | Y | Y |
| m69 | F | Y | F | F | Y | F | F | Y | F | F | Y | F | F | Y | F | F | Y | F |
| m70 | H | F | Y | H | F | Y | H | F | Y | H | Y | Y | H | F | Y | H | F | Y |
| m71 | Y | Y | F | Y | Y | F | Y | Y | F | Y | Y | F | Y | Y | F | Y | Y | F |
| m73 | F | H | H | F | H | H | F | H | H | F | H | H | F | H | H | F | H | H |
| m74 | H | F | F | H | F | F | L | H | Y | L | H | Y | L | F | F | Y | F | F |
| m76 | Y | F | F | Y | F | F | H | H | F | H | H | F | H | H | F | H | H | F |
| m77 | F | Y | Y | F | Y | Y | F | Y | Y | F | Y | Y | F | Y | Y | F | Y | Y |
| m78 | H | F | F | Y | H | F | H | F | F | Y | H | F | H | F | F | Y | H | F |
| m79 | F | Y | H | F | Y | H | F | Y | Y | H | H | Y | H | H | Y | H | H | Y |
| m81 | H | L | H | H | H | L | H | H | L | H | H | L | H | H | L | H | H | L |

**Figure S12. Mutant library constructed for the second round**

For the 27 m10 mutants in the library, the amino acids at $X_4$, $X_5$, and $X_6$ positions are summarized. One of the four expected amino acids was introduced for all positions.

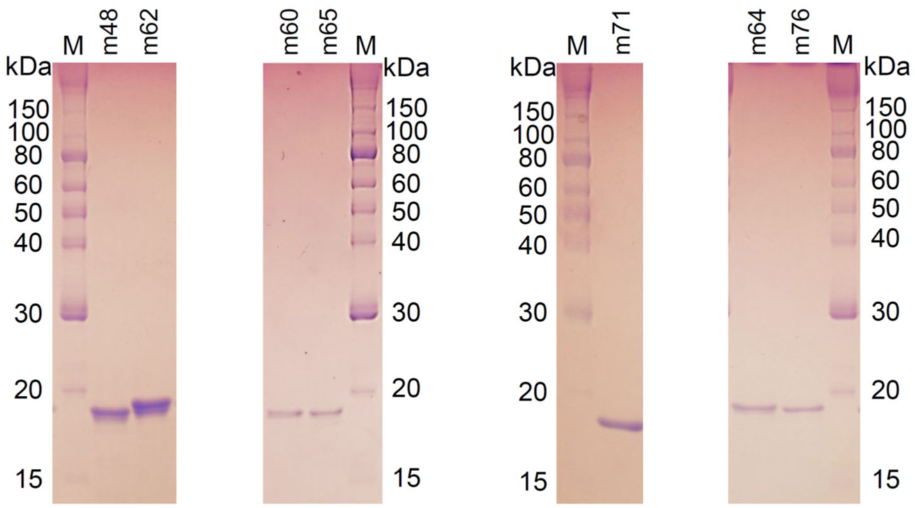


**Figure S13. SDS-PAGE images of the m10 mutants**

Lane M contains SIMASIMA Unstained Broad Range Protein Ladder (Cosmo Bio, Tokyo, Japan). No bands of contaminant proteins were observed, indicating that m48, m60, m62, m64, m65, m71, and m76 were of high purity.

**Table S1. Combinations of phosphorylated ssDNA oligos and primer sets for isothermal amplification.**

Underlined nucleotides represent the annealing sites of the reverse primers, and double-underlined nucleotides represent those of the forward primers. The RE ring was prepared using a split oligo DNA (ACCAGGAACACCATCACCACGACC) to connect two 5′-phosphorylated ssDNAs. N, D, Y, and W represent mixed bases (N: A, C, G, or T; D: G, A, or T; Y: C or T; and W: A or T).

| Template name | 5' phosphorylated ssDNA | Base length | Forward primer | Reverse primer |
|---|---|---|---|---|
| R | TCACCTTACTCAGGGCGCGGTGACTCTCCATACTCG<br>GGTCGTGGCGACAGCCCGTATAGCGGTCGTGGCGA<br>CAGTCCGTATTCTGGTCGTGGTGAT | 96 | GATATACATATG<br>AAAGGGCGCGGT<br>GACTCTCC | ATGATGAGTCGG<br>CCATGAGTAAGG<br>TGAATCACCACG<br>ACC |
| E | GGTGTTCCTGGTGTAGGTGTCCCAGGTGTCGGCGT<br>GCCGGGTGTGGGTGTACCAGGCGTTGGCGTACCGG<br>GCGTAGGGGTACCTGGTGTT | 90 | GATATACATATG<br>AAAGTAGGTGTC<br>CCAGGTGTCGG | ATGATGAGTCGG<br>CCAACCAGGAAC<br>ACCAACACCAGG<br>TAC |
| RE | TCACCTTACTCAGGGCGCGGTGACTCTCCATACTCG<br>GGTCGTGGCGACAGCCCGTATAGCGGTCGTGGCGA<br>CAGTCCGTATTCTGGTCGTGGTGATGGTGTTCCTGG<br>TGTAGGTGTCCCAGGTGTCGGCGTGCCGGGTGTGG<br>GTGTACCAGGCGTTGGCGTACCGGGCGTAGGGGTA<br>CCTGGTGTT | 186 | GATATACATATG<br>AAAGGGCGCGGT<br>GACTCTCC | ATGATGAGTCGG<br>CCATGAGTAAGG<br>TGAAACACCAGG<br>TAC |
| E-3NDT | GGTGTTCCTGGTGTAGGTGTCCCAGGTGTCGGCGT<br>GCCGGGTNDTGGTGTACCAGGCNDTGGCGTACCGG<br>GCNDTGGGGTACCTGGTGTT | 90 | GATATACATATG<br>AAAGTAGGTGTC<br>CCAGGTGTCGG | ATGATGAGTCGG<br>CCAACCAGGAAC<br>ACCAACACCAGG<br>TAC |
| m10-3YWY | GGTGTTCCTGGTCACGGTGTCCCAGGTATTGGCGTG<br>CCGGGTYWYGGTGTACCAGGCYWYGGCGTACCGG<br>GCYWYGGGGTACCTGGTCAC | 90 | GATATACATATG<br>AAACACGGTGTC<br>CCAGGTATTGG | ATGATGAGTCGG<br>CCAACCAGGAAC<br>ACCGTGACCAGG<br>TAC |
